# Coenzyme Q10 alleviates the mitochondrial damage by high-fat load in hepatocytes of spotted seabass (*Lateolabrax maculatus*) via promoting mitophagy

**DOI:** 10.1101/2025.03.07.642039

**Authors:** Yi-xiong Ke, Xiao-jiang Mao, Xue-shan Li, Ling Wang, Kai Song, Chun-xiao Zhang, Bei Huang, Kang-le Lu

## Abstract

Coenzyme Q10, as a natural fat-soluble compound, can play a role in protecting mitochondria, but the mechanism is still unclear. Here, we explored the mechanism of coenzyme Q10 enhancing mitochondrial function using hepatocytes of spotted seabass. Three groups were set: normal medium as control group, fatty acid group containing 100 μmol/L FA (FA group), and 100 μmol/L FA and 5 μmol/L coenzyme Q10 group (FA+COQ10). After the culture, the results showed that FA treatment significantly increased the triglyceride content in the cells. Bodipy staining showed that many lipid droplets appeared in the FA group, while coenzyme Q10 reduced triglycerides content and lipid droplets. Moreover, coenzyme Q10 significantly reduced the content of ROS in cells. After scavenging ROS, the liver cell damage caused by FA was alleviated, the mitochondrial membrane potential and its mitochondrial metabolic enzyme activity were restored, and the ATP content was increased. Further analysis showed that FA significantly down-regulated the expression of mitophagy key genes pink, parkin and lc3b, while up-regulated the expression of p62. Through mitochondrial fluorescence staining and mtDNA content detection, it was found that the number of mitochondria in FA-treated cells decreased significantly, while the number of mitochondria increased significantly after FA+COQ10 treatment. This indicates that coenzyme Q10 can significantly promote the mitophagy process. In order to further study whether the enhancement of mitochondrial function by coenzyme Q10 is related to the activation of autophagy, we set up FA group, FA+COQ10 group and FA+COQ10+Mdivi-1 group (pretreatment with mitophagy inhibitor Mdivi-1). After Mtphagy Dye staining, it was found that the number of autophagosomes in the FA+COQ10+Mdivi-1 group was lower than that in the FA+COQ10 group, indicating that the activation of mitophagy by coenzyme Q10 was inhibited. The results of this study indicate that coenzyme Q10 enhances mitochondrial function and alleviates excessive fat deposition dependent on PINK1-mediated mitophagy.

## 1. Introduction

In fish, the liver plays a key role in lipid metabolism[1]. When the source of fat is greater than the way, the liver is prone to excessive fat deposition and fatty liver occurs[2]. In the early stage of fatty liver, the lesion is not obvious but it will affect the liver function and weaken the resistance of the fish[2]. Previous studies of our laboratory have shown that the occurrence of fatty liver in fish is often accompanied by mitochondrial dysfunction [3, 4], a fact that has slso been in zebrafish, grass carp and yellow catfish.[5–7].

The specific manifestations include the decrease of key rate-limiting enzyme activity in the tricarboxylic acid cycle, the abnormal decrease of respiratory chain complex enzyme activity, the decrease of fatty acid β oxidation rate, the decrease of mitochondrial membrane potential, and the decrease of ATP production[4, 8, 9], which is similar to the results of non-alcoholic fatty liver in mammals[10, 11]. Mitochondria, as a highly dynamic organelle, play a key role in lipid metabolism and cell homeostasis[11]. Oxidative stress caused by excessive production of ROS caused by mitochondrial damage led to the second blow of fatty liver[12, 13]. Therefore, the increase in the number of damaged mitochondria may accelerate the progression of fatty liver disease, and the restoration of damaged mitochondria may be an effective way to alleviate fatty liver.

As a kind of cell-specific autophagy, mitophagy is essential for the removal of damaged mitochondria. When mitophagy is hindered, it may cause adverse effects such as damaged mitochondrial deposition, structural rupture, and oxidative stress[14]. These effects are often similar to the results of metabolic disorders caused by non-alcoholic fatty liver disease. Studies have found that mitophagy is closely related to liver fat metabolism, and abnormal autophagy may be involved in the “double blow” of metabolic liver disease[15]. In addition, mitophagy can remove triglyceride (TAG) to regulate lipid metabolism, thereby preventing liver steatosis[16]. Studies have shown that mitophagy can promote mitochondrial production, which further emphasizes its important role [17].

Coenzyme Q10, as a natural fat-soluble compound, it is widely distributed in the human body and generally exists in fish [18]. Coenzyme Q10 exists in most eukaryotic cells, especially in mitochondria. It has important functions such as scavenging free radicals, reducing oxidative stress damage, improving mitochondrial function, promoting oxidative phosphorylation, producing ATP and promoting energy metabolism[19, 20].

Coenzyme Q10 can play a role in protecting mitochondria, but the mechanism is still unclear. Here, we explored the mechanism of coenzyme Q10 enhancing mitochondrial function using hepatocytes of spotted seabass.

## 2. Materials and methods

### 2.1. Reagents

Coenzyme Q10 (C111044) was purchased from Aladdin (Shanghai, China). With reference to the existing research dissolution method [21], the coenzyme Q10 powder was dissolved in N, N-dimethylformamide(DMF, D112004, Aladdin, Shanghai, China) solvent under dark conditions to prepare a yellow transparent coenzyme Q10 solution with a concentration of 10 mmol/L.

Oleic acid (O1008) and palmitic acid (P0500) were purchased from Sigma company and dissolved in 20 % BSA made from PBS solution to prepare a stock solution with a final concentration of 10 mM. Mitophagy inhibitor Mdivi-1 (338967-87-6) was purchased from Glpbio (USA).

### 2.2. Cell culture

A strict ethical code for animal experiments was established by Jimei University’s Ethics Committee on Animal Experiments (JMU202403034).The primary hepatocytes were derived from spotted seabass livers using a tissue block culture method, as described in a previous study [22]. The basal medium was composed of 20 % feal bovine serum (FBS, Gibco) and penicillin 100U/m), streptomycin (0.1mg/mL) (Solarbio, Beijing, China) and DMEM/F12 (DMEN/F12, Shanghai Basal Media Technologies Co, Ltd, China).

### 2.3. Experiment I

In order to explore the effect of coenzyme Q10 on hepatocytes of spotted seabass, three experimental groups were set up, which were the control group (Control) without any treatment, the high-fat group (FA) treated with 0.1 mM FA, and the treatment group with 5μm COQ10 added to the medium of the FA group (FA + COQ10). The treatment time of each group was 12 h.

### 2.4. Experiment II

In order to study the physiological significance of activating mitophagy, according to the published literature[23], in the FA+COQ10 experimental group, after the cells adhered to the wall, 10 μM mitophagy inhibitor Mdivi-1 ( 338967-87-6, Glpbio) was first added for 2 h. The experimental group was named FA+COQ10+Mdivi-1.

### 2.5. Cell viability

After the liver cells were resuspended, the hepatocytes of spotted seabass were cultured in 96-well plates. After the cells were stably adherent, the medium containing different concentrations of drugs was replaced, and the culture was continued until the set test time. Subsequently, the medium containing 1% CCK-8 solution was replaced and incubated at 28 °C for 4 h. Finally, the absorbance was measured by a microplate reader at 450 nm, and the cell viability was calculated and analyzed according to the instruction formula.

### 2.6. Cell biochemical parameters

The hepatocytes of spotted bass were inoculated in a 6-well cell culture plate and cultured until the cells adhered stably. The cells were divided into three groups: control group, high-fat group with 100umFA and 100umFA + 5umCOQ group. After 12 hours of treatment, the cells were collected by trypsin digestion, and then the cells were washed twice with 1XPBS (pH = 7.2-7.4, P1020, Solarbio, Beijing, China) for the next measurement.

Commercial kits were used to detect triglyceride (TAG) (E1013, Applygen, Beijing, China), protein (PC0020, Solarbio, Beijing, China), aspartate aminotransferase (AST) (A010-2-1, NanjingJiancheng, Nanjing, China), alanineaminotransferase (ALT) (C009-21, NajingJian cheng, Nanjing, China), lactatedehydrogenase (LDH) (A020-2-2, NanjingJiancheng, Nanjin, China), citrate synthase (CS) (A108-1-2, Nanjing Jiancheng, Nanjing, China). In addition, blank control was set up in the determination of experimental indexes to eliminate background error.

Additionally, the activity of succinate dehydrogenase (SDH) (BC0955, Solarbio, Beijing, China) was determined by a commercial kit. After the cells were inoculated in a 6-well cell culture plate and subjected to experimental treatment, the cell culture medium was discarded, washed twice with PBS buffer, and then the kit extraction reagent was added. After ultrasonic lysis, the lysate was collected and centrifuged at 11, 000 g at 4 °C for 10 min. The supernatant was taken for determination. The microplate reader was preheated for 30 min and read at 600 nm wavelength for 20 s absorbance. It was quickly placed in a 37 °C water bath for 5 minutes and read for 5 minutes and 20 seconds absorbance. Finally, the succinate dehydrogenase activity of each group was calculated according to the formula.

The content of ATP was determined by enhanced ATP assay kit (S0027, Beyotime, Shanghai, China). After the liver cells were treated, the cells were collected and lysed with 200 μL of lysis buffer for 10 min. The lysis buffer was collected and centrifuged at 12, 000 g at 4 °C for 5 min, and the supernatant was taken for determination. Firstly, the ATP detection working liquid was configured according to the instructions, and 100 μL per hole was added to the all-black background enzyme plate, and placed at room temperature for 5 min to eliminate the background ATP, and then 20 μL of the sample to be tested was quickly added. The chemiluminescence detection was performed by the multi-function microplate reader (Varioskan LUX, Thermo Scientific), and the RLU value was substituted into the standard curve reading.

The content of reactive oxygen species ( ROS) ( S0033S, Beyotime, Shanghai, China) was determined by flow cytometry. The treated cells were digested with trypsin, and then incubated with a medium containing 10 μM DCFH-DA fluorescent probe ( S0033S, Beyotime) ( 28 °C, 20 min). After incubation, the cells were tested on the machine. The content of malondialdehyde (MDA) (A003-4-1, Nanjing Jiancheng, China) was determined by adding the cell lysate after the experimental treatment of the cells, and then the determination was carried out according to the kit instructions. Finally, the microplate reader was used to read at 532 nm. Similarly, the activity of superoxide dismutase(SOD) (A001-3-2, Nanjing Jiancheng, China) was also determined according to the kit instructions. After inoculation of hepatocytes (2 × 106 cells / well), the experimental treatment was carried out. After the experimental treatment, the IRPA lysate (R0030, Solarbio, Beijing, China) containing PMSF was added for lysis treatment, and the supernatant was collected by centrifugation for subsequent determination. In addition, blank control was set up in the determination of experimental indexes to eliminate background error.

### 2.7. Fluorescence Image

In order to evaluate the content of lipid droplets in cells, according to previous studies, Bodipy method was used for staining. Firstly, liver cells were inoculated on cell climbing slides, and treated with experiment I and II after cell adherence. Then, the cells were stained with BodipyTM 493/503 (D3922, Invitrogen) and DAPI at a concentration of 2μg/mL. In order to evaluate the mitochondrial content of liver cells, we seeded the cells in a glass-bottomed petri dish. After the treatment of experiment I, the cells were incubated with 200 nM Mitotracker Green probe (C1048, Beyotime) (28°C, 20 min); in order to evaluate the content of reactive oxygen species in the cells, we treated the cells according to experiments I, and incubated the cells with a medium containing 10 μM DCFH-DA fluorescent probe (S0033S, Beyotime) (28°C, 20 min); in order to evaluate the changes of mitochondrial membrane potential, we treated the cells according to experiment I, and incubated the cells with JC-1 fluorescent probe (C2006, Beyotime) (28°C, 20min); in order to evaluate the apoptosis of cells, we treated the cells according to experiment I, and incubated the cells with 500 μL Hoechst 33258 staining solution in the dark (room temperature, 10 min); in order to evaluate the co-localization of mitochondria and lysosomes, cells were treated with experiment I and incubated with 200 nM Mitotracker Green and 75 nM Lyso-Tracker Red (C1046, Beyotime) fluorescent dyes (28°C, 20 min); in order to evaluate the mitophagy of cells, cells were treated with experiment I, and cells were incubated with 100 nM Mtphagy Dye mitophagy probe (MD01, Dojindo) (28°C, 20 min). After incubation, the nuclei were stained with Hoechst 33342 (100X) (C1028, Beyotime) at a concentration of 5μg/mL (28°C, 20min). Finally, all fluorescence was observed and photographed using a Leica sp8 laser confocal microscope, and use image j software for subsequent image analysis.

### 2.8. Gene expression

The total RNA of liver was extracted by FastPure cell / tissue total RNA isolation kit (Vazyme Biotech Co., Ltd., China) from Nanjing Novazan Company, and the specific operation was carried out according to the instructions. The total RNA was immediately placed on ice after extraction, and then the purity and concentration of RNA were detected at 260/280 nm using a microspectrophotometer (NanoDrop 2000, Thermo Scientific). 1% agarose gel electrophoresis was performed to evaluate the integrity of the sample RNA. Finally, the total RNA of the liver was reversely transcribed using the kit (R211-01, Vazyme) provided by Nanjing Novisan to prepare cDNA, and the specific steps also follow the instructions.

The reverse-transcripted cDNA was first diluted proportionally with RNase-free ddH2O, and real-time fluorescence quantitative PCR was performed using Novozan’s SYBR ® Green I (Q711-02, Vazyme) kit and using a fluorescence quantitative PCR instrument (QuantStudio TM 6 Flex, Applied Biosystems). The CDS sequence of the target gene was derived from the transcriptome data of Lateolabrax maculatus[24].

After searching and comparing the gene sequence in the NCBI database, the primer sequence was designed by Primer 5.0 program, and then sent to Shanghai Shenggong Bioengineering Technology Co., Ltd.for primer synthesis. The real-time fluorescence quantitative reference genes in this experiment were all β-actin, and the relative expression of genes was calculated by 2^−ΔΔCt^ method. The primer sequences used in this experiment are shown in Table 1.

**Table 1.**
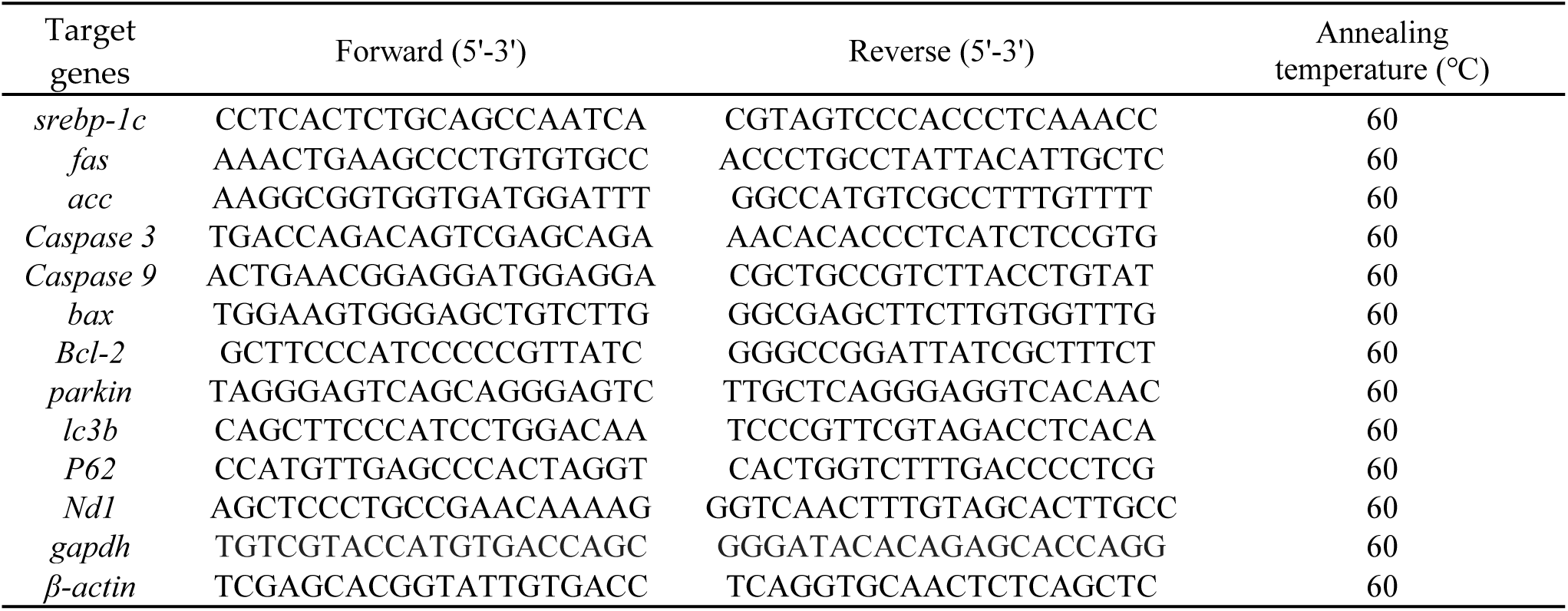
RT-qPCR primer sequence of *L. maculatus*.

### 2.9. Quantitative detection of mitochondrial mtDNA

The total DNA in the cells was isolated using the Nanjing Novozyme DNA extraction kit (DC102, Vazyme). The concentration and integrity of DNA were determined and tested by micro-spectrophotometer and agarose gel electrophoresis. Mitochondrial target gene ND1 (NADH dehydrogenase subunit 1) and internal reference gene GAPDH were designed by Primer Premier 5.0 software. The relevant primer sequence is detailed in Table 1, and the subsequent fluorescence quantitative PCR analysis method is the same as 2.8. Mitochondrial DNA copy number was analyzed by relative quantitative analysis, and the final results were expressed as the percentage of the control group.

### 2.10. Statistical Analysis

In this study, all data were analyzed by one-way analysis of variance (One-way ANOVA) using SPSS21.0 software. The differences between the experimental groups were compared by Tukey method. If *P* < 0.05, the difference was statistically significant. All results are expressed as mean ± standard error (Mean ± S.E.M.).

## 3. Results

### 3.1. Effects of coenzyme Q10 on fat deposition and metabolism

As shown in Figure 1, the addition of 5μM coenzyme Q10 in the FA medium could significantly reduce the fat deposition. As shown in Fig. 1B, adding 5μM coenzyme Q10, the number of lipid droplets decreased. As shown in Fig. 1C, D, compared with the control group, the FA group significantly up-regulated the expression of fat synthesis-related genes srebp1-c, fas and acc, and significantly down-regulated the expression of lipolysis genes atgl and hsl *(P* < 0.05). Compared with FA, coenzyme Q10 significantly down-regulated the expression of fas, acc and srebp-1 *(P* < 0.05), and up-regulated the expression of atgl and hsl *(P* < 0.05). At the same time, compared with the control group, pparα in the FA group was significantly down-regulated *(P* < 0.05), and cpt-1α had a downward trend but no significant difference *(P* > 0.05). Compared with the FA group, the pparα and cpt-1α genes in the coenzyme Q10 group were significantly up-regulated *(P* < 0.05).

**Figure 1.**
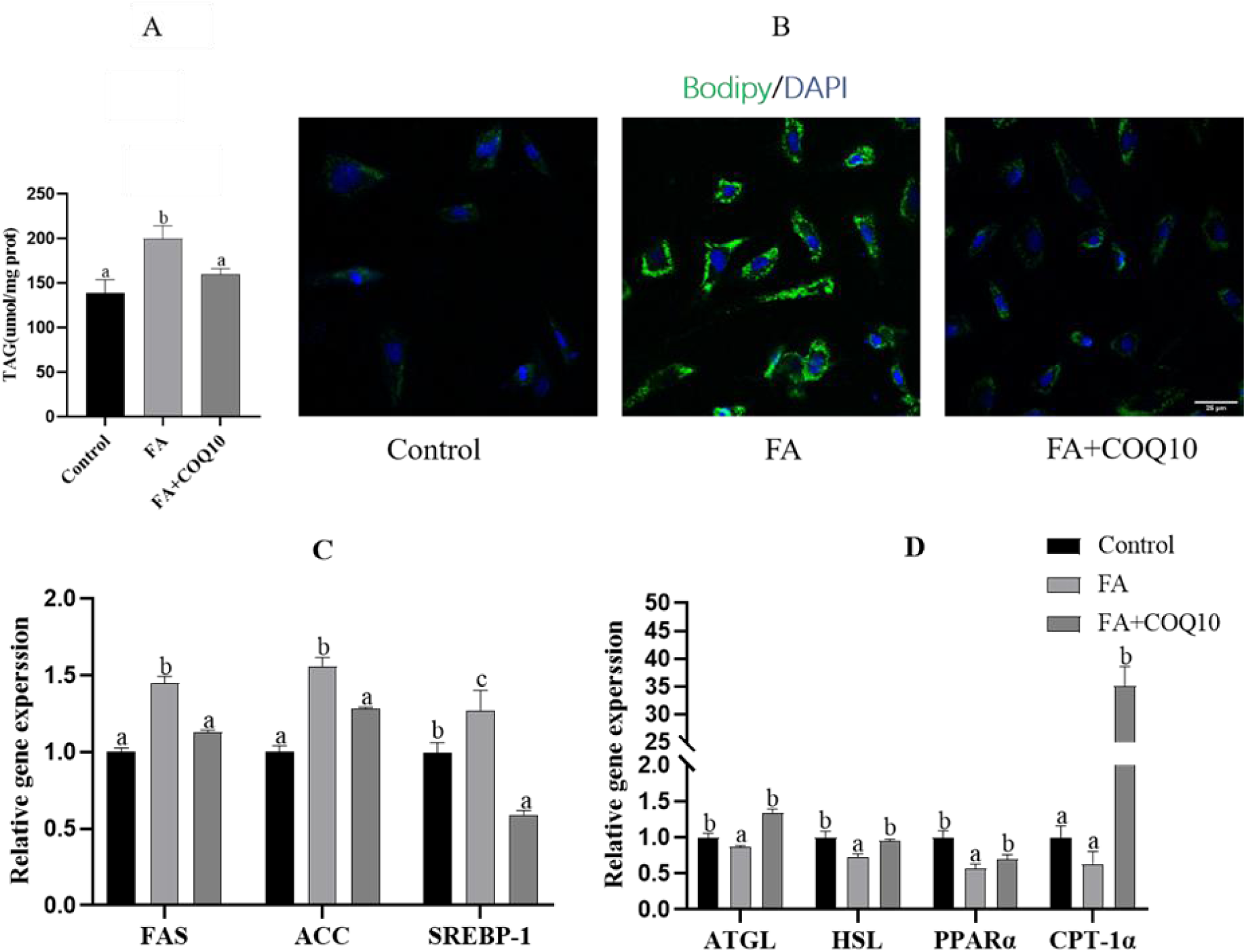
Effect of coenzyme Q10 on fat deposition in hepatocytes of L. maculatus. A: The triglyceride content of liver cells of L. maculatus was incubated with coenzyme Q10 for 12 h. B: Lipid droplets were stained with BODIPY493/503 (green) in the liver cells of L. maculatus. The nucleus was highlighted by Hoechst 33342 (blue) with a scale of 25 μm. C, D: mRNA levels of lipid metabolism-related genes (fas, acc, srebp-1, atgl, hsl, pparα, cpt1-α) in hepatocytes of L. maculatus. All values are exhibited as mean ± SE. The values with different superscripts (a, b, c) are significantly different at *P* < 0.05 (Tukey’ test).

### 3.2. Effect of coenzyme Q10 on ROS content and apoptosis

As shown in Fig. 2A, B, compared with the control group, the ROS in the FA group was significantly increased; after adding COQ10, the content of ROS decreased significantly *(P* < 0.05). Similarly, intracellular ROS fluorescence staining showed that the fluorescence intensity of ROS in the control group and COQ10 group was weak, and the fluorescence intensity of FA group was higher, indicating that high fat produced more ROS, while COQ10 could scavenge ROS. As shown in Fig.2C, with apoptosis-Hoechst staining, compared with the control group, the nucleus of the FA group showed dense staining and fragmented dense staining, indicating that apoptosis occurred, but the nucleus of the coenzyme Q10 group and the control group showed normal blue. FA group significantly up-regulated the expression levels of *bax*, *caspase3* and *caspase9 (P* < 0.05), and down-regulated the expression level of *bcl-2 (P* < 0.05). These results were reversed after the addition of coenzyme Q10 (Fig.2D).

**Figure 2.**
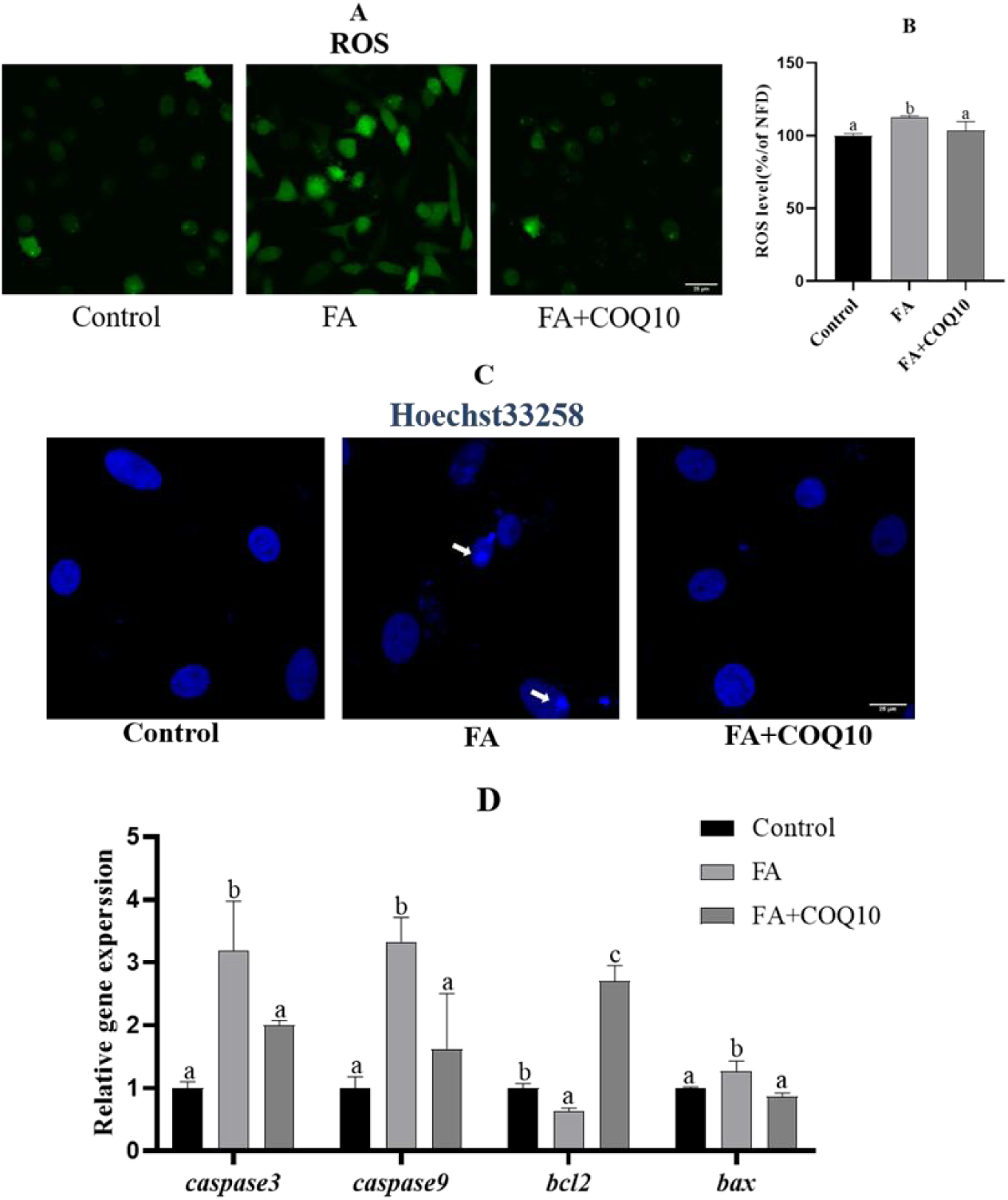
Effects of coenzyme Q10 on hepatocytes ROS content and apoptosis in L. maculatus. A: DCFH-DA dye was used for fluorescent staining of reactive oxygen species, ruler = 25 μm, B: flow cytometry to determine the relative content of ROS, DCFH-DA staining to detect ROS content, C: Hoechst33258 staining to detect apoptosis, scale = 25 μm, D: mRNA levels of apoptosis-related genes (caspase3, caspase9, bcl2, bax) in the liver of L. maculatus. All values are exhibited as mean ± SE. The values with different superscripts (a, b, c) are significantly different at *P* < 0.05 (Tukey’ test).

### 3.3. Effects of COQ10 on mitochondrial function

As shown in Figure 3A, B, C, compared with the control group, the CS and SDH activity and ATP content of the FA group were significantly decreased *(P* < 0.05), and the CS, SDH activity and ATP content of the coenzyme Q10 group were significantly increased *(P* < 0.05). The mitochondrial membrane potential JC-1 staining showed that the green fluorescence in the liver cells of the FA group increased and the red fluorescence decreased, indicating that the mitochondrial membrane potential was damaged at this time, but the green fluorescence decreased and the red fluorescence increased after the addition of coenzyme Q10, indicating that coenzyme Q10 can alleviate the damage of mitochondrial membrane potential *(P* < 0.05) (Fig.3D).

**Figure 3.**
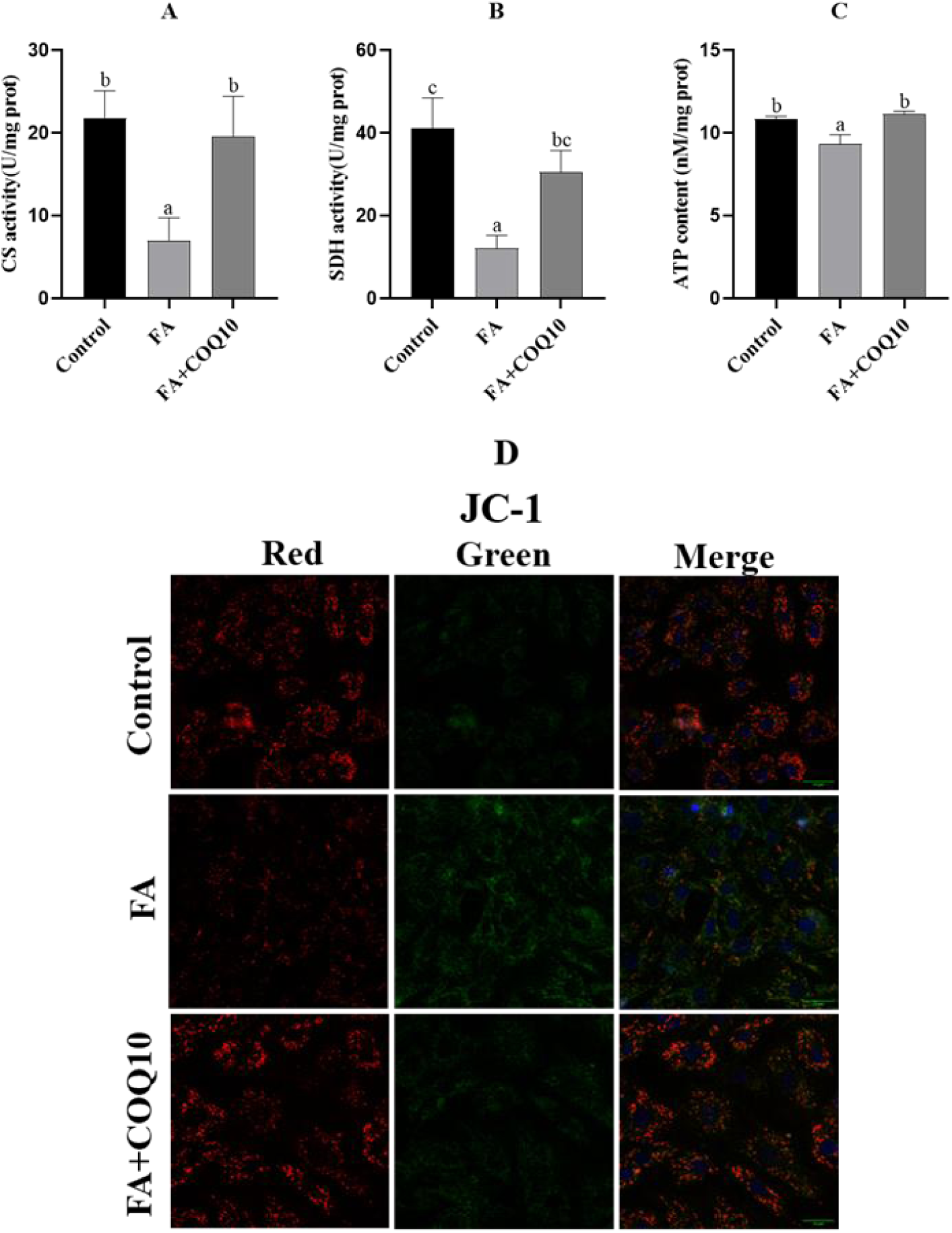
Effects of coenzyme Q10 on mitochondrial function in hepatocytes of L. maculatus. A: citrate synthase activity (CS), B: succinate dehydrogenase activity (SDH), C: ATP content, D: JC-1 staining was used to observe the changes of mitochondrial membrane potential in hepatocytes of L. maculatus. All values are exhibited as mean ± SE. The values with different superscripts (a, b, c) are significantly different at *P* < 0.05 (Tukey’ test).

### 3.4. Effects of coenzyme Q10 on mitochondrial biogenesis and autophagy

As shown in Fig. 4A, the relative content of mtDNA in each group was detected. Compared with the control group, the relative content of mtDNA in the FA group decreased significantly, and the content of mtDNA in the coenzyme Q10 group was restored *(P* < 0.05). The results of mitochondrial Mitrotracker green staining are shown in Fig.4B.The green fluorescence intensity of FA group was weak, indicating that the mitochondrial content decreased. The green fluorescence of the control group and the COQ10 group was stronger, indicating that the mitochondrial content of the cells was more. The fluorescence staining of mitophagy bodies is shown in Fig.4C.Compared with the control group, mitophagy in the FA group was inhibited, which was manifested by a significant decrease in the number of autophagosomes. This inhibition was alleviated after the supplementation of coenzyme Q10. As shown in Fig.4D, the fluorescence co-localization of mitochondria and lysosomes in the FA group was significantly weakened, and it was significantly enhanced after the addition of coenzyme Q10. As shown in Fig.4E, compared with the control group, the relative expression of mitophagy-related genes pink1, parkin and lc3b in the FA group was significantly down-regulated, and the relative expression of p62 gene was significantly up-regulated *(P* < 0.05). Coenzyme Q10 could significantly up-regulate the expression of these autophagy-related genes pink1, parkin and lc3b, and down-regulate the expression of p62 *(P* < 0.05).

**Figure 4.**
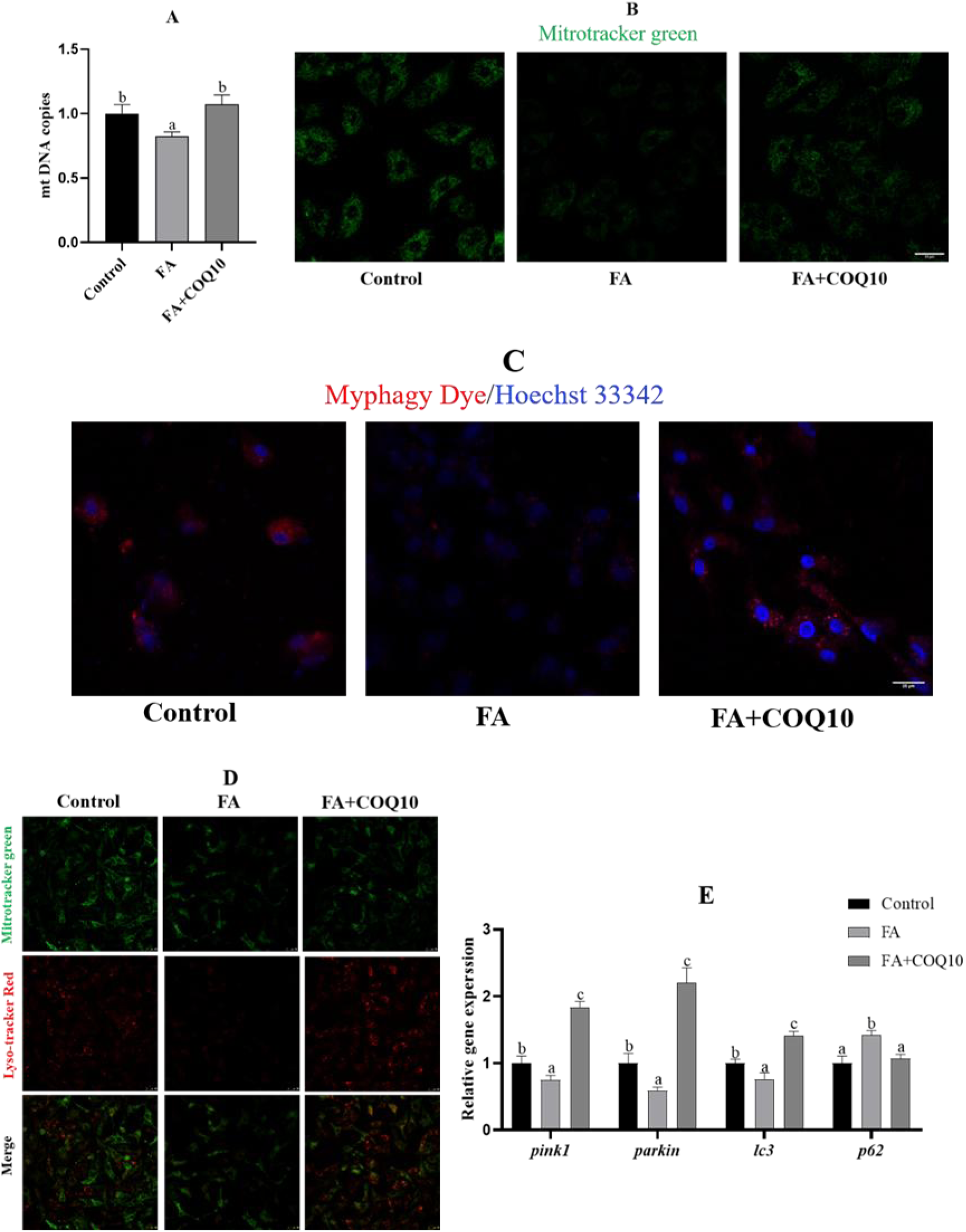
Effects of coenzyme Q10 on mitochondrial biogenesis and autophagy in hepatocytes of *L. maculatus*. A: the relative content of mtDNA, B: Mitrotracker green staining to observe the change of mitochondrial content, ruler = 25 μm. C: Myphagy Dye staining was used to observe the changes of autophagosomes. The nucleus was highlighted with Hoechst 33342 (blue), and the scale was 25 μm. D: Co-localization of mitochondria (green) and lysosomes (red) in hepatocytes of *L. maculatus*, ruler = 25 μm. E: mRNA levels of mitophagy-related genes (*pink1*, *parkin*, *lc3*, *p62*) in hepatocytes of *L. maculatus*. All values are exhibited as mean ± SE. The values with different superscripts (a, b, c) are significantly different at *P* < 0.05 (Tukey’ test).

### 3.5. Inhibition of mitophagy attenuated the effect of coenzyme Q10

As shown in Fig.5A, compared with the FA+COQ10 group, the number of autophagosomes in the Mdivi-1 group decreased; the results of Bodipy staining and triglyceride content showed that the triglyceride content and the number of lipid droplets in the Mdivi-1 group were significantly increased *(P* < 0.05) (Fig.5B, C). The activity of CS and SDH and the content of ATP in Mdivi-1 group were decreased *(P* < 0.05) (Fig.5D, E, F). At the same time, the effect of coenzyme Q10 on improving the activity of AST, ALT and LDH in the supernatant to alleviate cell damage was also inhibited by Mdivi-1 (Fig.5G, H, I).

**Figure 5.**
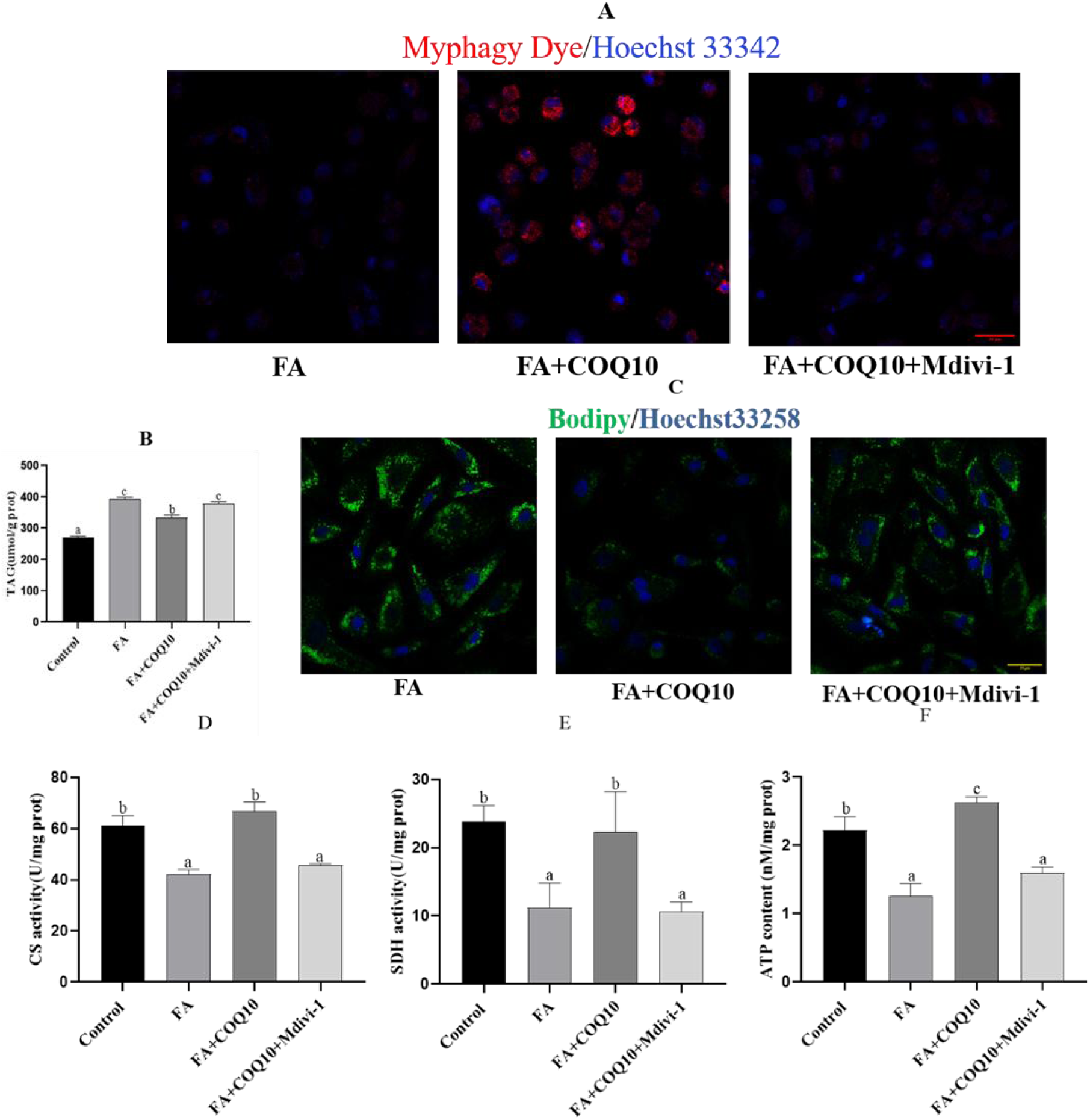
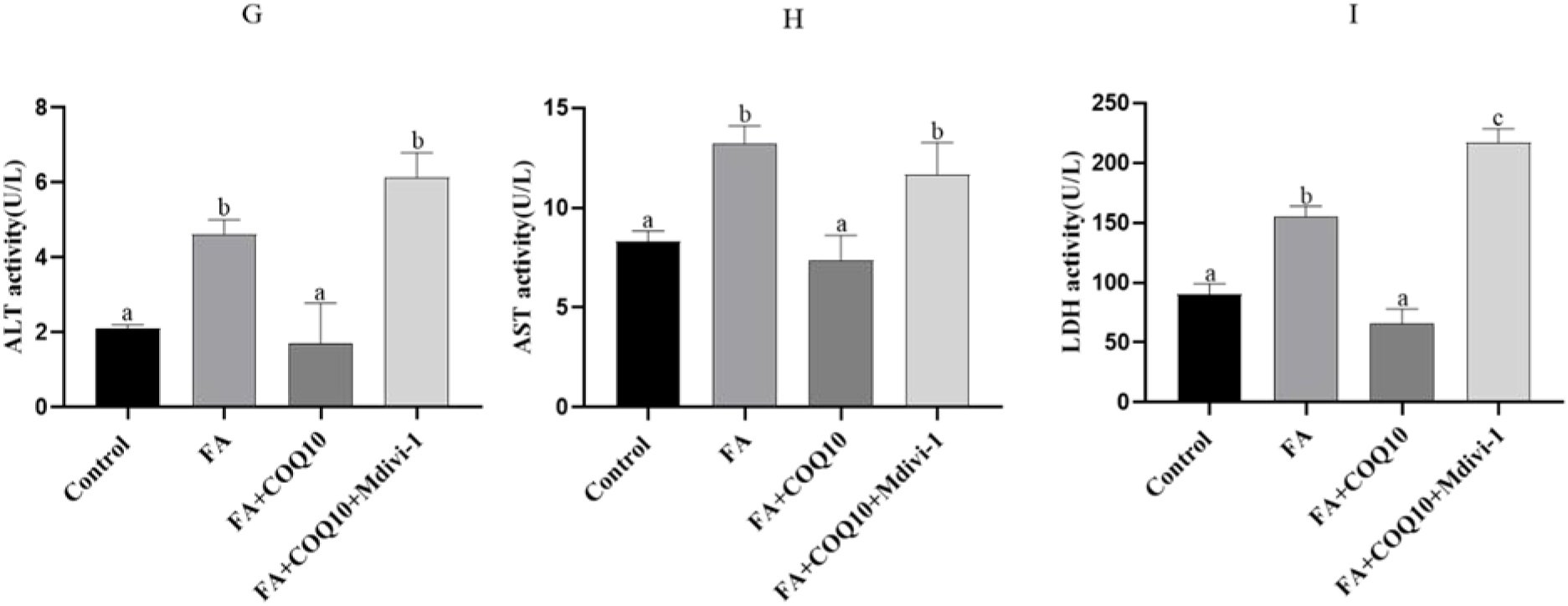
Inhibition of mitophagy by mitophagy inhibitor Mdivi-1 attenuated the effect of coenzyme Q10. A: After using the mitophagy inhibitor Mdivi-1, Myphagy Dye staining was used to observe the changes of autophagosomes. The nucleus was highlighted with Hoechst 33342 (blue), and the scale was 25μm. B, C: Triglyceride content in hepatocytes liver cells of spotted seabass, using BODIPY493/503 (green) staining lipid droplets. The nucleus was highlighted by Hoechst 33258 (blue) with a scale of 25 μm. D: citrate synthase activity (CS), E: succinate dehydrogenase activity (SDH), F: ATP content, G: alanine aminotransferase activity (ALT), H: aspartate aminotransferase activity (AST), I: lactate dehydrogenase (LDH) activity. All values are exhibited as mean ± SE. The values with different superscripts (a, b, c) are significantly different at *P* < 0.05 (Tukey’ test).

## 4. Discussion

As a lipid-soluble substance present on most eukaryotic biofilms, coenzyme Q10 is actively involved in the process of mitochondrial energy metabolism and is closely related to maintaining the normal function of mitochondria. In this study, 0.1 mM FA treatment could significantly promote the fat deposition of cells and 5 μm coenzyme Q10 effectively reduced fat deposition, Previous studies have found that coenzyme Q10 has no adverse effect on HepG2 cells at 0-100μm and has a good lipid-lowering effect[25].

The disorder of liver lipid metabolism may be caused by the formation of new fat, the activation of lipid uptake and the decrease of lipid catabolism. *Srebp-1* regulates the expression of many genes involved in adipogenesis, including *acc*, *fas* and *scd1* [26, 27]. The results showed that COQ10 inhibited the expression of Srebp-1, *fas* and *acc*, and inhibited the formation of new fat. *Atgl*, *hsl*, *pparα* and *cpt-1α* are key enzymes in lipolysis and fatty acid β-oxidation[28]. The results showed that coenzyme Q10 promoted the expression of these genes, which indicates coenzyme Q10 can promote the utilization of fatty acids.

Mitochondria are important organelles in the cell. Mitochondrial dysfunction is closely related to neurodegenerative diseases, myocardial and skeletal muscle abnormalities[29, 30].The most important thing is that mitochondria are closely related to metabolic diseases. Some researchers regard nonalcoholic fatty liver disease as a mitochondrial disease. Mitochondrial damage is a research focus in the study of NAFLD[31, 32].

The stability of mitochondrial structure and function is closely related to liver lipid metabolism. A large number of studies have shown that high-fat load can cause the damage of mitochondria[8, 9, 33, 34]. In this study, FA caused the decrease of CS and SDH activities and ATP content. Also, the JC-1 staining showed the decrease of mitochondrial membrane potential. Thus, FA affected mitochondrial energy metabolism and reduced ATP content.However, coenzyme Q10 reversed these results and restored mitochondrial membrane potential. These results demonstrate the positive effect of coenzyme Q10 on mitochondrial function. Studies have shown that COQ10 can enhance mitochondrial function, scavenge oxygen free radicals, and has a good preventive effect on metabolic diseases, and an important feature of metabolic diseases is the lack of coenzyme Q10, which also confirms that our supplementation of coenzyme Q10 can well alleviate mitochondrial damage and fat deposition in the liver of spotted bass[35, 36].

Mitochondrial membrane, as a barrier to protect mitochondria, will significantly increase its permeability when mitochondria are damaged, cytochrome C is released from mitochondria into the cytoplasm, and then transformed into a factor that promotes apoptosis, and activates the cascade response of *Caspase9/3* to trigger mitochondrial-controlled apoptosis [37]. *Bcl-2* and *bax*, members of the *bcl* protein family, play a key role in regulating the apoptosis[38]. The results of this experiment showed that FA up-regulated the expression of *caspase3/9* and promoted apoptosis. Coenzyme Q10 could down-regulate the expressions of *caspase3/9* and *bax*, and up-regulate the expression of *bcl-2*. In this study, coenzyme Q10 significantly reduced the degree of apoptosis. Moreover, FA caused oxidative stress in hepatocytes and abnormal accumulation of ROS. Coenzyme Q10 can reduce ROS content in cells and restore antioxidant capacity.

Previous studies have shown that mitophagy is the key to maintaining the renewal and regeneration of mitochondria[17]. This experiment explored whether the effect of coenzyme Q10 on enhancing mitochondrial function depends on the activation of mitophagy. Mitophagy Dye was used to stain mitophagy, and the number of autophagosomes in the FA group decreased significantly, while coenzyme Q10 significantly activated the mitophagy process. At the same time, Mt DNA content detection and mitochondrial fluorescence staining also found that the number of mitochondria was significantly reduced when mitophagy was inhibited, and the addition of coenzyme Q10 could promote the generation of mitochondria, which again showed that coenzyme Q10 could promote mitophagy and promote the generation of mitochondria[39, 40].

PINK1-mediated mitophagy is a classical mitophagy pathway. When mitochondrion is damage, PINK1 stabilizes on the outer membrane of mitochondria, activates Parkin ubiquitin ligase activity, recruits autophagy core protein LC3BII to form mitochondrial autophagosomes, and finally fuses with lysosomes to degrade damaged mitochondria[41, 42]. As an essential autophagy receptor in the process of selective autophagy (mitophagy), P62 acts as a bridge between polyubiquitinated substrates and autophagosomes and thus the accumulation of P62 indicates that mitophagy is hindered [43]. By co-localization staining of mitochondria and lysosomes, the formation of mitophagy lysosomes can be reflected to a certain extent. This study showed that the formation of mitophagy lysosomes in the FA group was inhibited, thereby preventing the removal of damaged mitochondria. On the contrary, coenzyme Q10 promoted the relative expression of pink1, parkin and *lc3b* and the formation of mitophagy lysosomes, and reduced the expression level of *p62* and increased the number of autophagosomes. These results indicate that coenzyme activates PINK1-mediated mitophagy. Furthermore, the fluorescence staining for mitochondria and lysosomes also revealed that FA blocked the autophagy process. In the COQ10 group, the number of lysosomes in hepatocytes was increased. Studies on muscle cells have also found that coenzyme Q10 can enhance the function of lysosomes to activate mitophagy[44], which indicates that coenzyme Q10 can activate mitophagy.

This study also used the mitophagy inhibitor Mdivi-1 to pretreat the FA+COQ10 group. Mtphagy Dye staining results showed that the activation of mitophagy was weakened after Mdivi-1 pre-treatment. The results of TAG content determination and Bodipy staining also showed that the lipid-lowering effect of coenzyme Q10 disappeared after Mdivi-1 pre-treatment. In addition, coenzyme Q10 increased the activities of CS and SDH, and the increase of ATP content was also reversed. These results confirmed that coenzyme Q10 activating mitophagy play key role in enhancing mitochondrial function and reducing fat deposition.

In summary, FA treatment caused abnormal accumulation of fat, inhibited mitochondrial formation and autophagy, and aggravated mitochondrial damage. Coenzyme Q10 can reduce fat deposition, reduce oxidative stress and enhance mitochondrial function. Further studies have shown that coenzyme Q10 can promote mitochondrial production and activate pink1-mediated mitophagy to remove damaged mitochondria, thereby improving mitochondrial function. Of course, in order to further determine the beneficial effect of coenzyme Q10 on spotted bass, in vivo experiments of spotted bass are needed to verify a series of results we explored on cells.

## Author Contributions

Yixiong Ke: Data curation, Writing – original draft. Xiaojiang Mao: Data curation, Formal analysis. Bei Huang: Methodology, Writing – review & editing. Chunxiao Zhang:Conceptualization, Project administration. Kai Song: Conceptualization, Resources. Xueshan Li: Methodology, Formal analysis. Ling Wang: Resources, Visualization. Kangle Lu: Funding acquisition, Supervision, Writing – review & editing.

